# Identification of proteins involved in *Trypanosoma brucei* DNA replication fork dynamics using nascent DNA proteomics

**DOI:** 10.1101/468660

**Authors:** Maria C Rocha-Granados, Yahaira Bermudez, Garvin Dodard, Anthula V Vandoros, Arthur Günzl, Michele M Klingbeil

## Abstract

DNA replication, transcription and chromatin remodeling are coordinated to ensure accurate duplication of genetic and epigenetic information. In regard to DNA replication, trypanosomatid parasites such as *Trypanosoma brucei* display unusual properties including significantly fewer origins of replication than model eukaryotes, a highly divergent Origin Replication Complex (ORC), and an apparent lack of several replication factor homologs. Although recent studies in *T. brucei* indicate functional links among DNA replication, transcription, and antigenic variation, the underlying mechanisms remain unknown. Here, we adapted an unbiased technology for the identification of replication fork proteins called iPOND (**<u>i</u>**solation of **<u>p</u>**roteins **<u>o</u>**n **<u>n</u>**ascent **<u>D</u>**NA) to *T. brucei*, its first application to a parasite system. This led to the mass spectrometric identification of core replication machinery and of proteins associated with transcription, chromatin organization, and DNA repair that were enriched in the vicinity of an unperturbed active replication fork. Of a total of 410 enriched proteins, among which DNA polymerase *α* and replication factor C were scoring in the top, around 25% of the proteins identified were of unknown function and, therefore, have the potential to be essential trypanosome-specific replication proteins. Initial characterization of a protein annotated as a Replication Factor C subunit (Tb927.10.7990), and a protein of unknown function (Tb927.3.5370) revealed that both proteins retain nuclear localization throughout the cell cycle. While Tb927.3.5370 appeared to be a dispensable gene, Tb927.10.7990 proved to be essential since its silencing caused a growth defect in procyclic cells, accumulation of zoids and impaired DNA replication. Future studies on the generated proteins list can contribute to the understanding of DNA replication dynamics in *T. brucei* and how replication is coordinated with other cellular processes to maintain genome integrity.

## Introduction

Eukaryotic DNA replication is strictly coordinated and regulated by numerous molecular machines to ensure genomic stability for future cell generations. DNA replication initiation is coordinated with cell cycle progression through the multiprotein Origin Recognition Complex (ORC) that plays an essential role by recruiting proteins that lead to the assembly of the replicative machinery with the assistance of regulatory components Cdc6 and Cdt1. The key factors <u>C</u>dc45, the <u>M</u>CM replicative helicase complex, and <u>G</u>INS proteins form the CMG complex that further recruits other replication factors such as the clamp loader Replication factor C (RFC), the clamp proliferating cell nuclear antigen (PCNA) and the three replicative DNA polymerases (*α*, *δ*, *ε*) leading to processive DNA replication [1,2].

Instead of the archetypical Origin Recognition Complex (Orc1-6) found in model eukaryotes, trypanosomes contain ORC1 and four other highly divergent ORC subunits (TbORC4, TbORC1b, Tb3120 and Tb7980) [3,4]. Components acting downstream of origin activation are conserved in trypanosomes including the MCM helicase, and portions of the DNA synthetic machinery [5]. However, initiation regulatory factors such as Cdc6 and Cdt1 are lacking, and Cdc45 displays an unorthodox mode of regulation being exported from the nucleus prior to mitosis [4]. Trypanosomes also lack some classical cell cycle checkpoints and it is also well known that there are significant differences in cell cycle regulation between different life cycle stages, although the molecular mechanisms underlying these differences remain elusive [6,7].

Additionally, eukaryotic DNA replication is characterized as having multiple origins that are often defined by local DNA structure and chromatin environment rather than by sequence determinants [8–11]. Chromatin environment influences the spatial and temporal distribution of DNA replication events that are also coordinated with transcription. In mouse embryonic cells, 85% of the origins of replication (ORIs) are associated with annotated transcriptional units. Additionally, ORIs with higher firing efficiency are located at CpG islands of promoters [12]. Furthermore, 50% of all activated human ORIs overlap with transcription start sites (TSS), suggesting co-linear progression of DNA replication forks and transcription complexes to prevent head on collisions and potential collapse of the replication fork [13].

Evidence for functional interplay between DNA replication and transcription was also found in the trypanosomatid organisms *Leishmania major* and *T. brucei*. In *L. major*, ORIs were mapped preferentially at transcription termination sites (TTS), genomic locations where RNA pol II is expected to slow or stall [14].

This study concluded that there is coupling between origin activity and transcription, where DNA replication opportunistically initiates from genomic regions that have been available for RNA pol II elongation [14]. A global analysis of DNA replication initiation in *T. brucei* revealed that transcription and DNA replication initiation are coordinated in terms of genomic position [15,16] In *T. brucei*, genes are arranged as polycistronic transcription units also known as directional gene clusters (DGCs). Transcription of DGCs initiates at multiple positions either in divergent strand switch regions (dSSRs) in which DGCs are arranged head-to-head or, in some cases, between two arrays that face the same direction in a head-to-tail region (HT) [17,18]. dSSRs and HTs are open chromatin regions occupied by acetylated histone H4 (H4K10ac), trimethylated H3 (H3K4me3), histone variants H2A.Z and H2BV, and bromodomain factor 3 [19–22]. The replication initiator protein, TbORC1 prefers the same epigenetic landscape binding to divergent DGCs and the junctions between HT units proximal to H4K10Ac marks. This strong association reveled an unprecedented level of functional interaction between transcription and DNA replication in a eukaryotic genome. How these two machineries are coordinated spatially and temporally remains an open question. Collisions between the replication and transcription machinery are likely more frequent given the large tracks of co-directionally transcribed genes suggesting coordination between the transcription and DNA replication machinery or robust DNA replication restart machinery to overcome fork collapse.

Replication fork dynamics in *T. brucei* have mainly been gene-by-gene investigations that limit studying the machinery in a temporal fashion. The current minimal list of DNA replication factors were identified using biased approaches such as sequence similarity searches based on known replication factors in other model organisms, and affinity purification of associated factors with an already known protein as bait [3,4,23–27] While these studies provided valuable information indicating that the DNA replication machinery differs substantially from its host counterpart, the inventory of DNA replication factors, especially of regulatory factors and trypanosomatid-specific factors, remains incomplete [26].

One technique used to analyze replication fork dynamics and identify the proteins at the replication fork is iPOND (**<u>i</u>**solation of **<u>p</u>**roteins **<u>o</u>**n **<u>n</u>**ascent **<u>D</u>**NA) [28], a method developed in the human system. In iPOND, newly replicated DNA is labeled with the thymidine analog 5-ethynyl-2’-deoxyuridine (EdU) [29]. EdU contains an alkyne functional group that enables the cycloaddition of a biotin azide. This click chemistry reaction [30] yields a stable covalent linkage, facilitating streptavidin capture of cross-linked biotinylated DNA-protein complexes [31]. Combination of the iPOND technology with mass spectrometry (MS) provides a site-specific analysis of replisomes by the identification of the proteins associated to replication forks and helps to determine how these factors are coordinated at the replication site to maintain the genome integrity [32]. iPOND offers an unparalleled ability to identify factors at active and damaged replication forks and to follow the spatial and temporal dynamics of these processes by varying the labeling period with EdU. Additionally, variations using pulse-chase experiments can distinguish between proteins that are specific to replication forks as opposed to proteins that are part of bulk chromatin. Recently, iPOND was adapted in vaccinia virus, the prototype poxvirus, to identify proteins involved in viral DNA replication [33]. In addition to known viral replication proteins, viral DNA-dependent RNA polymerase and transcription initiation and elongation factors were identified on nascent DNA. This suggested that there is temporal coupling of DNA replication and transcription at active replication forks in poxviruses [33].

In this study we adapted and applied iPOND technology for the first time in a human-pathogenic parasite, *T. brucei*. By coupling iPOND with label-free mass spectrometric quantification using iBAQ (Intensity-based absolute quantification) [34], we were able to identify 410 proteins that were cross-linked and enriched with nascent DNA. The list includes known replication factors together with DNA repair, transcription, splicing, and chromatin organization factors. Nearly 25% of the data set were proteins of unknown function. Overall, we obtained a panoramic view of the cellular processes that appear to be coordinated with DNA replication and might help to maintain genomic stability. Additionally, we selected two proteins for initial characterization. These are a putative Replication Factor C subunit (RFC) and a protein of unknown function. Both proteins displayed nuclear localization. Only RFC proved to be essential in PCF cells. Cells in which this protein was depleted exhibited a DNA replication defect and growth impairment.

## Materials and Methods

For Primer sequences refer to Supplemental Table 1 (S1 Table).

**Table 1.** Comparison of mammalian and *T. brucei* DNA replication parameters

| Parameter | Mammalian cells<br>(293T)<sup>a</sup> | *Trypanosoma brucei*<br>(procyclic form) |
| --- | --- | --- |
| Genome size | 3000 Mbp<sup>b</sup> | 26 Mbp |
| % of cells in S phase | 50%<sup>c</sup> | 20%-30%<sup>d</sup> |
| Replication time | 480 min<sup>e</sup> | 90 min |
| Replication rate | 1.5 kb/min<sup>f</sup> | 3.7 kb/min |
| ~Kbp labeled in 10 min | 15 | 37 |
| Estimate firing forks | 2138<sup>g</sup> | 78<sup>h</sup> |
| Kbp labeled/fork | 32070 | 2886 |
| Number of cells | 1.6 x 10<sup>9</sup> | 3 x 10<sup>10</sup> |
| Cells in S phase | 8 x 10<sup>8</sup> | 7.5 x 10<sup>9</sup> |
| Total labeled kbp | 2.56 x 10<sup>13</sup> | 2.16 x 10<sup>13</sup> |
| Total µg labeled DNA | **28.1 µg** | **23.7 µg** |
| Total µg of DNA | 5399.1 | 854.6 |
| % DNA labeled | 0.52% | 2.77% |

<sup>a</sup>Human embryonic kidney cells [31]
<sup>b</sup>[58]
<sup>c</sup>[59]
<sup>d</sup>[56]
<sup>e</sup>[60]
<sup>f</sup>[61]
<sup>g</sup>[62]
<sup>h</sup>See calculations in S1 Appendix

### Plasmid construction

(i) <u>PTP tagging</u>. For PTP allelic tagging of *TbORC1, Tb427.03.5370* and *Tb427.10.7990* the C-terminal coding sequence was PCR amplified from *T. brucei* 427 genomic DNA and ligated into either pC-PTP-NEO [35] or pC-PTP-PURO [36]. In pORC1-PTP-NEO, p5370-PTP-NEO and p7990-PTP-PURO the corresponding gene sequences comprised the 3’ terminal 1098, 699, and 498 bp of the coding regions, respectively. All final constructs were sequenced. For genome integration, pORC1-PTP-NEO was linearized with SalI, pTb5370-PTP-NEO with AvaI, and pTb7990-PURO with XcmI.

(ii) <u>RNAi</u>. A pStL (stem-loop) vector for inducible gene silencing of each iPOND candidate was constructed as previously described [37]. Briefly, 542 bp or 417 bp of Tb427.03.5370 or Tb427.10.7990 coding sequence respectively were PCR amplified from *T. brucei* 427 genomic DNA using appropriate primers (S1 Table). Final pStL5370 and pStl7990 vectors were linearized with EcoRV for genome integration.

### *Trypanosoma brucei* Cell Lines

Cultivation of the PCF *Trypanosoma brucei brucei* Lister 427 strain [38] and of the 29-13 strain for conditional gene silencing [37] was carried out as described. To generate cell lines expressing PTP-tagged proteins or inducible RNAi, 7.5 μg of linearized plasmid was transfected by nucleofection using an Amaxa nucleofector 2b (Lonza) [39] and cells were selected with either G418, puromycin or phleomycin. In the cell line ORC1^PTP/WT^, the second *TbORC1* allele was deleted using the ApaI/NotI fragment from the pKOORC1-Hyg plasmid and selected using hygromycin [40]. The concentrations of selecting drugs in medium were 50 μg/ml of G418, 1 μg/ml of puromycin, 40 μg/ml of hygromycin and 2.5 μg/ml of phleomycin. Transgenic cell lines were cultured for no more than three weeks. For each transfection, correct DNA integration was confirmed by PCR of genomic DNA with at least one oligonucleotide hybridizing outside of the transfected nucleotide sequence (S1 Table). Conditional gene silencing experiments were performed by incubating trypanosomes in medium containing 1 μg/ml of tetracycline as described [37]. The single expresser clonal cell line ORC1^PTP/KO^ P2C2 (ORC1SE) was used for all iPOND experiments (10 hour doubling time).

### *Trypanosoma brucei* iPOND

The original Cortez iPOND method was followed with several modifications [28]. A total of 3 • 10^10^ log phase TbORC1PTP cells were labeled with 150 μM EdU (Santa Cruz) for 10 min. This length of time should label approximately 37 kb of DNA (see calculations in S1 Appendix). In the pulse-chase experiment, EdU labeling was followed by washing of cells once with temperature-equilibrated medium containing 150 μM thymidine and incubation of cells in medium containing 150 μM thymidine for a 60 min chase period prior to pelleting and cross-linking. Cells were pelleted at 3,000 x *g* for 7 min, immediately resuspended at approximately 7.5 μ 10^8^ cells/ml in SDM-79 medium containing 1.1% formaldehyde, and incubated for 20 min at room temperature (RT) to cross-link DNA-protein complexes. The reaction was quenched by adding 2 M glycine to a final concentration of 0.125 M and incubated for 5 min at RT. Fixed samples were washed three times with cold 1X phosphate buffer saline (PBS), and cell pellets flash frozen and stored at −80°C. For further processing, samples were thawed on ice for 30 min, resuspended with 0.25% Triton-X 100 in PBS in a volume to give an estimated cell density of 1 • 10^9^ cells/mL homogenized with 5 passes using a glass dounce, and then incubated for 30 min at RT with gently shaking. Samples were washed once with PBS containing 0.5% BSA and once with PBS, and the pellet was resuspended in click reaction cocktail (0.2 M biotin-azide, 50 mM sodium ascorbate, 10 mM CuSO_4_) at a volume of 5 ml for every 1 • 10^10^ total cells, and rotated for 2 hours at RT. Following the click reaction, cells were washed as specified above and resuspended in SME buffer (0.25 M sucrose, 10 mM MOPS pH 7.2, 2 mM EDTA, 1 mM μMSF, 1μg/ml leupeptin and one tablet of cOmplete EDTA-free Protease Inhibitor Cocktail (Roche)) containing 0.1% NP40 (cell density of 1 • 10^9^ cells/mL). This suspension was homogenized 30 times with a glass dounce and incubated for 1 hour on ice with gentle shaking prior to nitrogen cavitation at 2250 psi (20 min equilibration in SME buffer with 0.1% SDS). If necessary, this step was repeated (1500 psi and 10 min of equilibration) until >85% cell lysis (via microscopy) was achieved. Samples were centrifuged at 1250 x *g* for 10 min to obtain an enriched nuclear fraction (P1, S1 Fig), washed (SME, then PBS) and resuspended in sonication buffer (0.7% wt/vol SDS in 50 mM Tris pH 8.0, 1 μg/ml leupeptin and one tablet of cOmplete EDTA-free Protease Inhibitor Cocktail) by dounce homogenization. Samples were sonicated 5X (10 min, 30s ^ON/OFF^) using the Bioruptor UCD-200 (Diagenode), and lysates cleared at 16,000 x *g* for 10 min at 4°C (Input). To capture biotinylated DNA-protein complexes, the Input sample was diluted with 1 volume of PBS, mixed with PBS-equilibrated Streptavidin-agarose beads (250 μl beads slurry) (Novagen), and incubated overnight 4°C (16-20 hours) using a rotator. Beads were pelleted (1,800 x *g*, 3 min, RT) and washed once with streptavidin wash buffer (2% wt/vol SDS, 150 mM NaCl, 50 mM Tris-HCl, pH 7.4), once with 1M NaCl, and twice with streptavidin wash buffer. Each wash step included 5 min of rotation and centrifugation (1,800 x g, 1 min, RT). Tubes were changed between washes. Proteins were eluted by incubating the beads at 95°C for 25 min in 2X Laemmli sample buffer. As negative controls, cells were not exposed to EdU but treated with only DMSO, and cells exposed to EdU but treated with the click chemistry cocktail that lacked biotin-azide (C^-^) were generated. Three biological replicates were performed for each condition except the pulse-chase experiment where only one experiment was analyzed.

### DNA Fragmentation Analysis

To analyze the extent of DNA fragmentation after sonication, lysate aliquots (50 μl) were subjected to cross-link reversal by adding NaCl to a final concentration of 0.2 M and incubating the samples overnight at 64 °C. Subsequently, samples were treated with 10 μl RNase A (20 mg/ml) for 30 min at 37 °C followed by Proteinase K (Ambion) treatment for 2 hours at 45 °C (20 μl of 0.5 M EDTA, 40 ul of Tris pH 6.7 and 10 ul of Proteinase K). Finally, DNA was prepared by phenol/chloroform extraction and ethanol precipitation, and quantified using a Nanodrop 8000 (Thermo Scientific). DNA samples (100 ng) were separated on a 1.5% agarose gel, stained with ethidium bromide and visualized under UV light.

### Dot Blot Click Chemistry Efficiency Test

Biotinylated tubulin oligonucleotides (standard curve) and sonicated input fraction DNA (2 μg per replicate) were treated with 1 M NaOH, heated at 55 °C (30 min) followed by addition of 2 M ammonium acetate to remove crosslinks. Oligo DNA and sonicated DNA samples (~100 ng) were spotted onto a nylon membrane (GE healthcare science). Oligo DNA (serially diluted ranging from 0.25 pmol to 16 pmol) was spotted in triplicate and input samples (~100 ng) were spotted in duplicate. Membrane was air-dried and cross-linked (Stratalinker 1800) then immediately placed in blocking solution (20% non-fat milk) for 2 hours at 37°C. Membrane was then rinsed three times in PBS + 0.1% Tween-20 and incubated with Avidin-HRP (1:3000, Life Technologies) for 30 min (37°C) followed by 3 washes in PBS + 0.1 % Tween-20, each for 15 min. ImageQuant LAS 4000 mini (GE healthcare science) was used for chemiluminescence detection and data were quantified using ImageJ (version 1.51S). DNA from input fractions of each iPOND condition were quantified by nanodrop.

### EdU and PTP Immunofluorescence

EdU incorporation for a 10 min pulse was confirmed using the Picolyl azide Toolkit (Life Technologies). Cells were labeled with 150 μM EdU for 10 min, immediately harvested, washed with ice-cold PBS and adhered to poly-L-lysine coated slides (5 min). Cells were then fixed in 3% paraformaldehyde (5 min, RT), washed in PBS containing 0.1 M μlycine (pH7.4) three times (5 min, RT), and permeabilized with 0.1% Triton X~100 in PBS (5 min, RT). After three additional washes with PBS (5 min, RT), Click chemistry was performed using the Click-iT Plus Alexa Fluor Picolyl Azide Toolkit (Life Technologies) according to manufacturer’s directions. Following click incubation for 1 hour at RT, cells were washed three times (5 min, PBS) and processed for immunofluorescence. Cells were incubated with anti-protein A antibody (Sigma) diluted 1:20,000 in PBS/1% BSA for 60 min, washed three times in PBS/0.1% Tween 20, and incubated with Alexa Fluor 594 goat anti-rabbit antibody diluted 1:250 in PBS/1% BSA for 60 min. DNA was stained with 3 μg/ml 4’-6’-diamidino-2-phenylindole (DAPI), and slides were washed 3 times in PBS prior to mounting in Vectashield (Vector Laboratories). Parasites were visualized and images captured either with a Nikon Eclipse E600 microscope with a cooled CCD Spot-digital camera (Diagnostic Instruments) using a 100X Plan Fluo 1.3 (oil) objective or with a Nikon N-SIM E superresolution microscope equipped with an RCA-Flash 4.0 sCOS camera (Hamamatsu Photonics) and a CFI SR Apochromate TIRF 100X (NA1.49) objective. Z-stacks (6 μM, 240 nm thickness) were acquired using the NIS-Elements Ar software. Image slices were reconstructed using default software parameters and 3D deconvolution using the automatic method in NSIM modality was applied. Images’ brightness and contrast were adjusted using Adobe Photoshop CS4 for presentation in figures.

### EdU labeling Quantification

(i) <u>Cell Cycle Analysis</u>. Cells were incubated with 150 μM EdU for 30 min, immediately harvested and processed as described above for EdU immunofluorescence. Cells were then incubated with rat monoclonal antibody YL1/2 (Abcam) (60 min, 1:500 in PBS + 1% BSA), washed three times in PBS + 0.1% Tween 20 and then incubated with Alexa Fluor 594 goat anti-rat, stained with DAPI and mounted as described above. At least 325 cells were scored for each time point. Only intact cells identified by phase contrast were included in the analysis. Cells were classified as EdU+ if fluorescence was detected. Position within the cell cycle was determined based on the kDNA morphology (DAPI staining) and number of basal bodies (YL1/2 staining). YL1/2 recognizes RP2, a protein that localizes to transitional fibers and is therefore a marker only for mature basal bodies [41,42]

(ii) <u>EdU Fluorescence Intensity</u>. Images were acquired using the Nikon E600/Spot digital camera system (described above). Non-saturating exposure times were used and non-adjusted images were analyzed using CellProfiler 3.0.0 [43] to measure EdU pixel intensity. Images were segmented with a DAPI signal to generate masks matching cell nuclei from which the mean EdU signal was calculated. A minimum of 220 EdU positive (EdU+) cells were analyzed from each time point. Data were represented using Prism 7 (GraphPad).

### SDS-PAGE and Western Blot Analysis

Samples were fractionated by SDS-PAGE and transferred overnight to PVDF membrane. For Histone 3 (H3) detection, membranes were blocked in PBS + 3% non-fat milk for two hours and incubated with rabbit H3 antiserum (1:50,000, 0.3% blocking solution) for 2 hours. For mono-, di- and trimethylated H3 at lysine 76 (H3K76) detection, membranes were blocked in PBS + 3% BSA for 2 hours, and incubated with antiserum in 0.3% blocking (H3K76me1, 1:300; H3K76me2, 1:1500; H3K76me3, 1:3000) for 2 hours. All H3 antibodies were kindly provided by Christian Janzen. ORC1PTP was detected using peroxidase-anti-peroxidase (PAP) (Sigma, 1:2000), and mtHsp70 was detected using *Crithidia fasciculata* specific antibody (1:10,000, gift from Paul Englund). Following primary antibody incubation, membranes were washed three times (PBS + 0.1% Tween-20) prior to incubation with appropriate horseradish peroxidase conjugated secondary antibodies in corresponding blocking solutions. Signal was detected with Clarity^TM^ ECL Blotting Substrate (NEB) using GE Imagequant LAS 4000.

### RNA Isolation and Quantitative reverse transcription-PCR Analysis

Total RNA was isolated from 5 • 10^7^ cells with Trizol (Thermo Fisher) according to the manufacturer’s specifications. RNA concentration was determined using Nanodrop, and 100 ng of total RNA was converted to cDNA using the High-Capacity cDNA Reverse Transcription Kit with RNase inhibitor and random primers (Thermo Fisher). qPCR was performed using QuantiNova SYBR Green PCR (Qiagen) with 1 μg of cDNA and 0.2 μM of primer (S1 Table) per reaction with a Stratagene MxPro 3000x thermocycler. All reactions were performed in triplicate. Gene knockdown was normalized with RNA of telomerase reverse transcriptase (TERT) [44].

### Mass Spectrometry Sample Preparation and Analysis

Final protein elutions (30 μl) of iPOND experiments were separated by SDS-PAGE. The gel of a complete sample lane was excised in sections and sent for liquid chromatography/tandem mass spectrometry (LC/MS/MS) analysis. Samples were further processed by the Keck Biotechnology Center of Yale University using their standard protocols. Briefly, each sample was resuspended in 25 mM ammonium bicarbonate containing 2.5 ng/μl digestion grade trypsin (Promega) and incubated at 37°C for 14 hours. After digestion, peptides were extracted from gels with two volumes of 80% acetonitrile, 0.1% formic acid for 15 minutes, then dried by speed vacuum. Peptides were dissolved in 30 μl of MS loading buffer (2% acetonitrile, 0.2% trifluoroacetic acid), with 5 μl injected for mass spectrometric analysis. LC/MS/MS acquisition was performed on a Thermo Scientific Q Exactive Plus coupled to a Waters nanoAcquity UPLC system.

For database searching, tandem mass spectra were extracted by Proteome Discoverer version 2.1.1.21 (Thermo Fisher). Charge state deconvolution and deisotoping were not performed. Data were searched in-house using the Mascot algorithm (version 2.6.0) (Matrix Science, London, UK). Mascot was set up to search a *Trypanosoma brucei* database (version 27, containing Lister strain 427 and TREU927, downloaded from http://tritrypdb.org/tritrypdb/). Search parameters used were trypsin digestion (strict) with up to 2 missed cleavages, peptide mass tolerance of 10 ppm, MS/MS fragment tolerance of 0.02 Da, and variable modifications of methionine oxidation, propionamide adduct to cysteine, and deamidation of asparagine and glutamine.

Scaffold (version Scaffold_4.8.2, Proteome Software Inc.) was used to validate MS/MS based peptide and protein identifications. Peptide identifications were accepted if they could be established at greater than 95.0% probability by the Scaffold Local FDR algorithm. Protein identifications were accepted if they could be established at greater than 99.0% probability and contained at least 2 identified peptides. Protein probabilities were assigned by the Protein Prophet algorithm (Nesvizhskii, 2003). Proteins that contained similar peptides and could not be differentiated based on MS/MS analysis alone were grouped to satisfy the principles of parsimony. Proteins sharing significant peptide sequence were grouped into clusters and were inspected individually.

### MS quantification

Protein abundance was estimated using the iBAQ (Intensity-based absolute quantification) value of each protein hit in the four different samples in the Scaffold program. The iBAQ value is based on the sum of all identified peptides intensities matching to a specific protein divided by the number of theoretically peptides observable, yielding an accurate proxy of protein abundance [34]. Proteins were considered to be enriched when they were identified in at least two of the three biological replicates and fulfilled the following criteria: (i) the protein hit must have a fold change over the No Click control that is equal or higher than 10, (ii) it must have a fold change over the DMSO control that is equal or higher than 1.5, and (iii) the protein should be a nuclear protein as determined by a recent nuclear proteome analysis [46], by the trypanosome genome wide localization resource [47], or have a predicted gene ontology (GO) term associated with a known nuclear protein. The lower fold change values for the DMSO control samples were likely due to the excess biotin azide from the click chemistry step that was incorporated as cofactor for some proteins (S2A Table).

To include proteins in our list that were identified in our nascent DNA analysis but absent in either negative control, e.g. obtained an infinite value (INF) for fold change, we set the fold change value from INF to the highest fold change value obtained in a particular experiment. To estimate a total score in order to rank the protein list the fold change (FC) EdU/No click and the FC EdU/DMSO were standardized to a 0-100 scale in each MS replicate. The average from each FC was calculated and the total score was determined by adding average FC EdU/No click by average FC EdU/DMSO.

### Bioinformatic Analysis

UniProt IDs of the proteins identified were acquired using the TriTrypDB database accession numbers [48] (http://www.tritrypdb.org/). However, each protein did not have a UniProt ID. GO term analysis and enrichment was performed using PANTHER classification system [49]. The UniProt IDs list was mapped against the *Trypanosoma brucei* reference list in PANTHER version 13.0 (released version 20171205) with the following selections: analysis type, overrepresentation test; annotation database, PANTHER GO-slim Biological process; and test type, Fisher’s Exact test with False discover rate (FDR) < 0.05. The results were sorted by hierarchically order to observe the enriched functional classes. Analysis of protein interaction networks was performed using STRING database [50]. Only interactions from curated databases and text-mining information were considered (confidence interaction score >0.65). The network was visualized using Cytoscape (version 3.6.0) and interaction groups were manually labeled based on GO term biological process [51].

Protein sequences from selected candidates were analyzed using the NCBI conserved domain database search (CDD) [52] to identify possible functional domains present in proteins of unknown function. Sequences were aligned using Clustal Omega with default parameters [53]. Motif search for Tb5370 was performed using the Eukaryotic Linear Motif (ELM) resource for Functional Sites in Proteins [54] with a cutoff for motif probability of 100.

## Results and Discussion

### *T. brucei* DNA replication properties for defining iPOND conditions

To purify proteins that associate with nascent DNA in *T. brucei*, we adapted the original iPOND method developed in the laboratory of Dr. David Cortez (Vanderbilt University) [32]. A major limitation of iPOND is the large amount of starting material needed to recover enough protein for proteomic analysis. Therefore, several aspects of trypanosome biology were considered when calculating the amount of starting material needed for efficient purification. These parameters included the *T. brucei* genome size (26 Mbp) [5], the duration of S phase (90 min) [55], the percentage of S phase cells in an unsynchronized population (20%-30%) [56], the estimated number of early firing origins, and the PCF *T. brucei* DNA replication rate (3.7 kb/min) [57]. To obtain an approximate equal amount of EdU labeled DNA with that in mammalian iPOND (2.56 • 10^13^ kbp corresponding to 28.1 μg DNA), we calculated that 3 • 10^10^ PCF cells were needed (2.59 • 10^13^ kbp) to obtain a comparable amount of labeled DNA (Table 1).

To establish iPOND in *T. brucei*, it was necessary to monitor the preparation of nuclei and the enrichment of nuclear proteins. Since there were no suitable antibodies for this purpose available, we took advantage of a previously established cell line in which the DNA replication initiation protein ORC1 was expressed from one allele with a C-terminal PTP tag [40]. To optimize analysis of ORC1-PTP we further manipulated the cells by knocking out the remaining wild-type allele, thereby generating a single expresser cell line (ORC1SE). We confirmed that the ORC1-PTP protein localized to the nucleus during all cell cycle stages and did not impact the fitness of the cells (S1A Fig). The ORC1SE cell line was used for all iPOND experiments and nuclear enrichment was tracked detecting the tag (S1D Fig).

### iPOND Optimization in *T. brucei*

Establishing the iPOND method in *T. brucei* required the optimization of several steps including EdU incorporation, click chemistry reaction, cell lysis, and DNA fragmentation (Fig 1A). Compared to mammalian cells where 10 μM EdU is sufficient to label DNA in a 10 min pulse, *T. brucei* PCF required 150 μM EdU to detect EdU-labeled DNA by fluorescence microscopy in the same period (Fig 1B). The lack of high affinity thymidine transporters can explain the higher concentration of EdU required for *T. brucei* labeling [63]. The high EdU concentration in our study is in accordance with other *T. brucei* labeling studies in which 100-300 μM EdU was applied for 1 hr to uniformly label the nucleus for studies on PCNA [24,25]. Our conditions, however, resulted in discrete spots that likely represent replication foci (Fig 1B).

**Figure 1.**
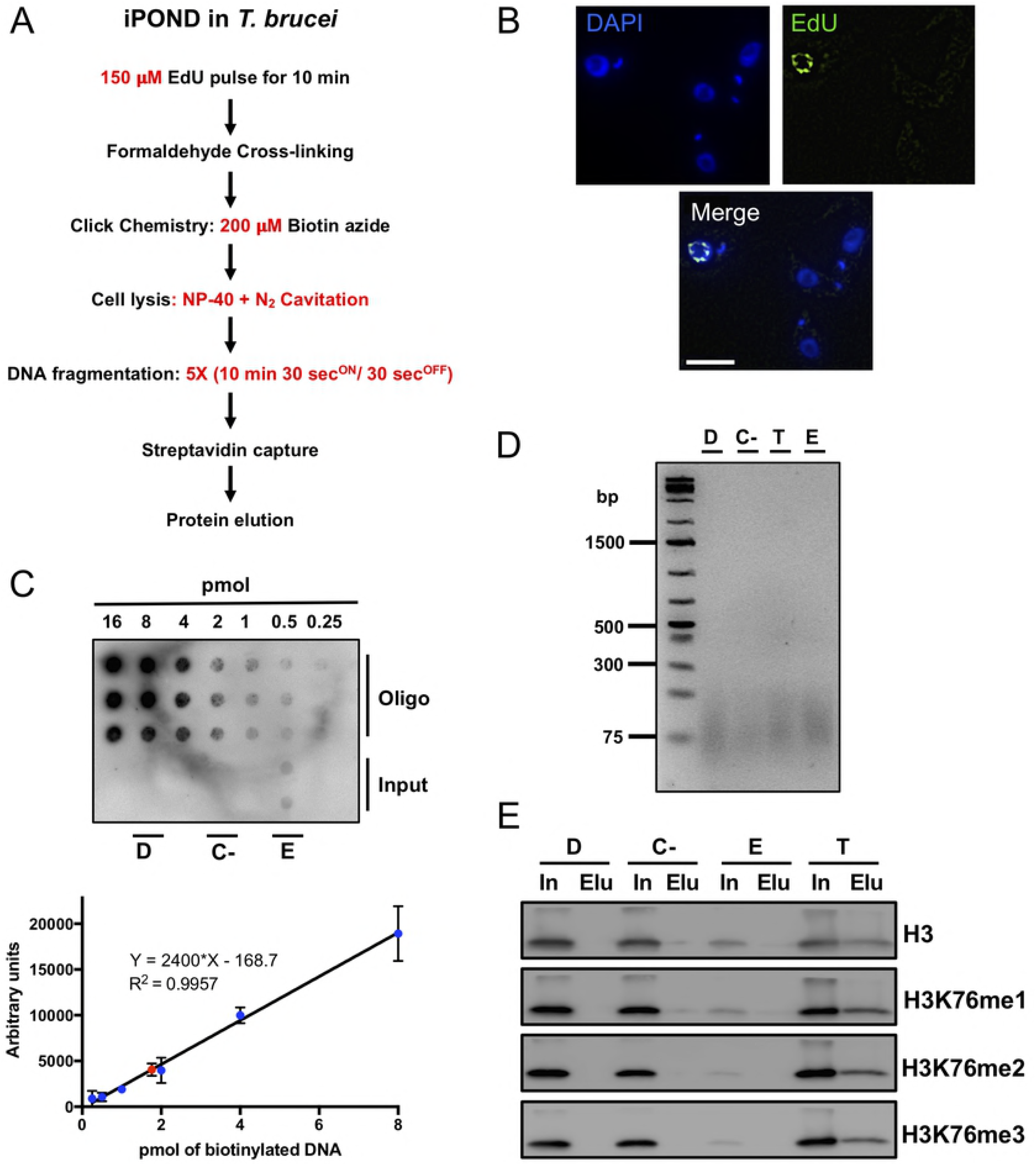
Optimization of iPOND in *T. brucei*. **(A)** Schematic overview of modified iPOND procedure for *T. brucei*. Red, modifications that were implemented compared to the original iPOND protocol performed in mammalian cells. **(B)** EdU incorporation in newly synthesized DNA in *T. brucei*. In a 10 min EdU pulse, newly synthesized DNA was successfully labeled. DNA is stained with DAPI. Size bar 5 □m. **(C)** Quantification of biotinylated DNA. Biotin incorporation was measured in sonicated samples (input) from negative controls (DMSO; D and Click-; C-), ThD chase (T), and EdU pulse (E) samples. Approximately 2 □g of DNA from sonicated samples (input) were applied to a Hybond membrane. Biotinylated DNA was detected using Avidin HRP. Graph, standards (blue circles) and EdU sample (red square) were plotted. **(D)** DNA fragmentation analysis. 5 cycles of sonication yielded DNA fragments between 50 bp to 200 bp, 100 bp fragments are enriched in all iPOND conditions. **(E)** Detection of Histone 3. H3 was used as a DNA-bound protein marker. Input (In) and final elution (Elu) fractions were probed against H3 and one, two and tree methylation groups present H3 Lysine 76 (H3K76met) in the four iPOND conditions.

The iPOND technology relies on click chemistry, which is a copper-catalyzed reaction that allows the cycloaddition of an alkyne functional group (present in EdU) to an azide (conjugated to biotin) yielding a stable covalent bond [64,65]. Based on the higher EdU concentrations for labeling, we optimized the efficiency of the click chemistry reaction in *T. brucei* by increasing the final concentration of biotin azide (200 μM) in the reaction cocktail 20-fold compared to mammalian conditions (10 μM). An increased amount of biotin azide was also used for iPOND in mouse embryonic cells [66].

Following the cross-linking step, standard lysis conditions resulted in incomplete lysis possibly due to a tightly cross-linked microtubule cytoskeleton [67]. Therefore, nitrogen cavitation was used in combination with detergent treatment to more efficiently lyse the cells (85-100% lysis). Despite the use of detergent in this extra step, we obtained a fraction enriched in seemingly intact nuclei (P1) with little visible contamination of kinetoplasts (mitochondrial DNA in *T. brucei*) or flagella (S1B, C Fig). Accordingly, ORC1-PTP was detected only in the P1 fraction while a mitochondrial matrix marker, Hsp70, was detected in both fractions (S1D Fig).

Compared to mammalian iPOND, additional rounds of sonication were required to shear the DNA to fragment sizes of 50-200 bp (Fig 1D), a range recommended for streptavidin capture of the cross-linked DNA-protein complexes. In order to monitor the amount of EdU-labeled DNA from sheared DNA samples (Input), we used a sensitive dot blot assay with a standard curve of biotinylated tubulin oligomers. For EdU pulse (E) experiments, 1.76 μMoles (~0.12 μg of DNA) of biotinylated DNA was detected from the 2 μg of DNA that was spotted, while there was no detection in the negative controls (Fig 1C). Biotinylated DNA was also detected in the thymidine chase experiment (S1E Fig). These modifications were critical to achieve approximately 24 μg of EdU-labeled DNA in a 10 min EdU pulse, comparable to the 28 μg regularly obtained in mammalian iPOND. In a 10 min pulse, we should label ~37 kbp. (Table 1). See S1 Appendix for details on calculations for iPOND.

### Validation of iPOND

To differentiate between proteins associated with nascent DNA, such as DNA polymerase *α* and the bulk of chromatin, such as modified histones, we compared iPOND from a 10 min EdU pulse with iPOND from a 10 min EdU pulse followed by a 60 min ThD chase. The short pulse should restrict DNA labeling to the vicinity of the replication fork, while during the ThD chase labeled DNA should have moved away from the fork and undergone chromatin deposition and remodeling. Histone deposition (without modification) is coupled with DNA replication, although the precise timing of deposition is still debated [68,69].

In *T. brucei*, localization of posttranslational-modified histones have been studied during the cell cycle [70,71]. For example, immunolocalization of mono-(H3K76me1) and di-methylated H3 variants (H3K76me2) indicated that these modified histones are detected during mitosis and cytokinesis but not during S phase [70], whereas the H3K76me3 modification occurs during all cell cycle stages [71].

To test whether unmodified or K76-methylated histone H3 was present in our final iPOND eluates we carried out western blot analysis using *T. brucei* specific immune sera. There is minor detection of H3 in the EdU sample while its signal increases in the ThD chase sample (Fig 1E, S2 Fig). Methylated H3 is mainly detected in the ThD chase sample (Fig. 1E). For some EdU samples minor amounts of H3K76me1 can be detected (Fig. 1E), however methylated H3 was rarely detected in the EdU samples (S2 Fig).

Quantification of band intensities revealed a 1.5% recovery of total H3 signal in the EdU elution compared to its input signal. In contrast, 13% of the H3 signal was recovered in the chase elution compared to the input. Additionally, the H3K76 methylation variants are enriched in the chase sample with an average of 8.5% recovery. H3K76me3 is undetectable in the EdU pulse sample even when increased cell equivalents were loaded and longer exposures analyzed (S2 Fig). Even though H3K76me3 was previously detected in all cell cycle stages, its deposition may not occur on newly replicated DNA, possibly explaining why we could not detect it in our EdU elution (E) (Fig. 1E). These data suggest that in procyclic *T. brucei* deposition of unmodified histones occurs on nascent DNA and posttranslational modifications occur as the DNA moves away from the replication fork. In accordance with this notion, Histone 4 lysine 4 acetylation (H4K4ac) was found to be cell cycle regulated with unmodified H4K4 being highest during S phase [72].

Based on these results, it appeared that the differences between the EdU and ThD chase iPOND samples likely represent early replicating conditions for the short pulse and matured chromatin for the chase conditions and, therefore, were suitable for proteomic analyses.

### Identification of proteins associated with nascent DNA

Proteins associated with nascent DNA (EdU pulse) from three biological replicates were isolated from gels, trypsin-digested, and analyzed by LC/MS/MS. To minimize the identification of false positives, we employed iBAQ to calculate fold changes (FC) between EdU pulse, negative controls and ThD chase samples (see methods for details). In addition, we restricted our analysis to known or putative nuclear proteins and disregarded proteins that are known standard contaminants of proteomic analyses such as low scoring ribosomal proteins, chaperones and proteins of retrotransposal origin [73]. A final score was calculated taking both the FC EdU/DMSO and the FC EdU/No Click into account (see S2 Appendix for details).

Based on these robust criteria, a total of 410 proteins were found to be enriched on nascent DNA (S2A Table). The genes of 98 proteins were annotated as “hypothetical conserved”, encoding proteins of unknown function (S2B Table). Gene Ontology (GO) enrichment analysis using the tool PANTHER [49] revealed 23 GO terms with >3-fold enrichment and a P-value of <0.001 (Fig 2A, S3 Table). The most abundant types of proteins were those involved in chromatin organization (fold enrichment, 10.7), transcription (9.91), DNA replication (7.43) and pre-mRNA splicing (7.03). To gain additional insight into the proteins enriched on trypanosome nascent DNA, we examined their potential relationships using the STRING database. In a STRING analysis, interactions are derived from multiple sources including curated databases that include known experimental interactions and text mining that incorporates prediction of interactions based on statistical links between proteins [50]. The analysis revealed a network of 9 clusters with abundant interactions between DNA replication and the DNA repair and nucleic acid metabolism clusters. However, there were also abundant links between DNA replication and transcription clusters (Fig 2B).

**Figure 2.**
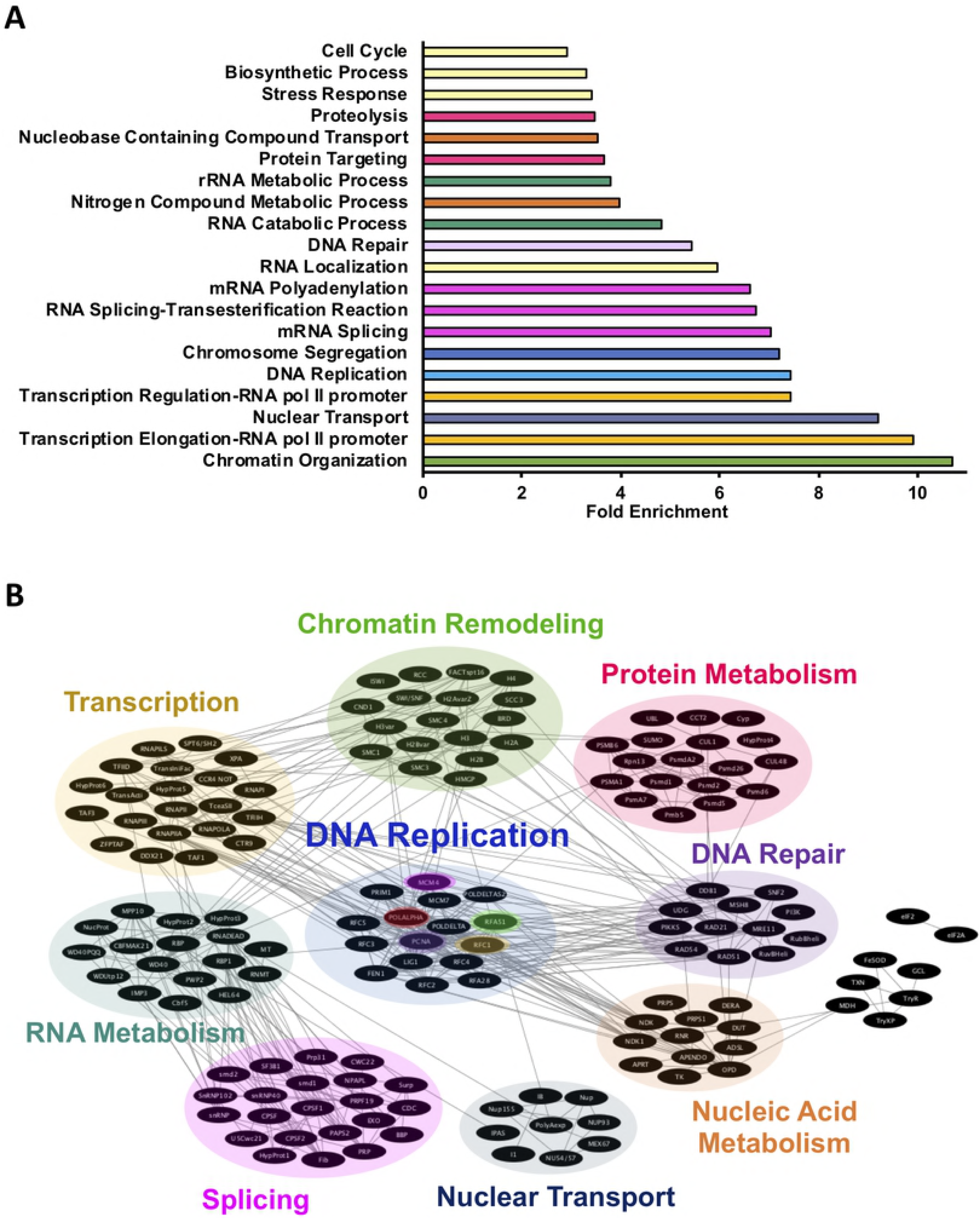
Gene ontology enrichment and protein network in *T. brucei* iPOND. **(A)** PANTHER Gene ontology (GO) analysis of the proteins identified on nascent DNA. Graph represents the fold enrichment of each GO term in a hierarchy order. Only GO terms with a fold enrichment above 3 were plotted. **(B)** Panoramic landscape of the protein-protein interaction network analysis of an active replication fork in *T. brucei* defined by STRING. Relevant interactions and most representative groups are displayed. The topology is organized according to functional classification. Highlighted nodes: DNA polymerase *α*, red; Replication Factor C, yellow; Replication Factor A subunit 1, green; MCM4, pink; and PCNA, purple.

As expected, known DNA replication proteins were enriched based on GO term analysis (fold enrichment of 7.43; P-value, 1.4 • 10^-9^) (Fig 2A, Table 2, S3 Table). Proteins forming the STRING replication cluster included MCM4 and MCM7 of the heterohexameric MCM complex (replicative helicase), DNA polymerases *α* (Pol *α*) and *δ* (Pol *δ*) (DNA synthesis), PCNA, replication factor C subunits (processivity), and FEN-1 endonuclease (Okazaki fragment processing). While several homologs of known *T. brucei* replication proteins were identified through iPOND and label free quantification in other systems [32,66], our analysis identified factors that were missing from these studies such as MCM and primase subunits (Table 2).

**Table 2.** DNA replication proteins identified in *T. brucei* iPOND

| **Tb427 Protein ID** | **Tb927 Protein ID** | **Product Description** | **Mol. Weight [kDa]** | **Ranking Position** | **Total Score** | **Identified by Cortez lab<sup>a</sup>** | **Identified by Ernforns lab<sup>b</sup>** |
| --- | --- | --- | --- | --- | --- | --- | --- |
| Tb427.08.4880 | Tb927.8.4880 | DNA polymerase alpha catalytic subunit | 152 | 9 | 134.02 | no | yes |
| Tb427tmp.01.4070 | Tb927.11.12250 | DNA replication licensing factor MCM4 | 93 | 13 | 132.20 | no | yes |
| Tb427.10.7990 | Tb927.10.7990 | ATPase, putative, replication factor 3 | 39 | 15 | 114.68 | no | yes |
| Tb427.03.830 | Tb927.3.830 | flap endonuclease-1 (FEN-1), putative | 44 | 71 | 77.62 | no | yes |
| Tb427tmp.01.7810 | Tb927.11.16140 | DNA replication licensing factor MCM7 | 81 | 93 | 73.57 | no | yes |
| Tb427.03.1130 | Tb927.3.1130 | DNA polymerase delta subunit 2, putative | 62 | 134 | 70.40 | no | yes |
| Tb427tmp.01.1310 | Tb927.11.9550 | replication factor C, subunit 4, putative | 38 | 149 | 69.20 | yes | yes |
| Tb427tmp.211.3310 | Tb927.9.12300 | replication factor C, subunit 3, putative | 40 | 180 | 51.86 | no | yes |
| Tb427.02.1800 | Tb927.2.1800 | DNA polymerase delta catalytic subunit, putative | 117 | 189 | 48.39 | yes | yes |
| Tb427.06.3890 | Tb927.6.3890 | replication factor C, subunit 2, putative | 39 | 216 | 42.93 | yes | yes |
| Tb427.07.2310 | Tb927.7.2310 | DNA primase small subunit, putative | 48 | 219 | 42.80 | no | no |
| Tb427tmp.02.3360 | Tb927.11.5650 | replication factor C, subunit 1, putative | 65 | 238 | 41.01 | yes | yes |
| Tb427.04.1330 | Tb927.4.1330 | DNA topoisomerase IB, large subunit | 79 | 283 | 37.39 | yes | no |
| Tb427tmp.160.3710 | Tb927.9.5190 | proliferative cell nuclear antigen (PCNA), putative | 32 | 311 | 28.61 | yes | yes |
| Tb427.06.4780 | Tb927.6.4780 | DNA ligase I, putative | 83 | 319 | 18.60 | yes | yes |
| Tb427tmp.01.0870 | Tb927.11.9130 | Replication factor A protein 1 | 52 | 321 | 17.37 | yes | no |
| Tb427.05.1700 | Tb927.5.1700 | replication Factor A 28 kDa subunit, putative | 28 | 388 | 7.89 | no | no |

### DNA replication proteins

PCNA, a key component of DNA replication as the sliding clamp, serves as a binding scaffold for numerous replication and DNA damage proteins [74]. In the replication cluster, PCNA was a key node to other DNA replication proteins but also to other clusters (Fig. 2B, purple node). Its interactions with a subunit of the replication factor complex (RFC1; Fig 2B, yellow node), and from RPA (RFA1; Fig 2B, green node), as well as interactions with MCM4 (Fig 2B, pink node) and DNA polymerase *α* (Pol *α*; Fig. 2B, red node) were expected. RFC1 is part of the clamp loader involved in PCNA loading, and RFA1 is part of the single-stranded DNA-binding protein complex RPA [75]. *T. brucei* RFC1 and RFA1 were enriched in our data set having ranking positions of 238 and 321 respectively, while PCNA ranked 311 (Table 2, S2A Table). *T. brucei* PCNA shows nuclear localization during the G1/S transition and S phase, regulation of its proper levels is critical for DNA replication and proliferation and is uniquely regulated by the kinase TbERK8 [24,25,27]. MCM4 ranked much higher at 13. The *T. brucei* MCM complex has been characterized and the single MCM4 subunit was able to unwind circular DNA *in vitro* as well as the complex [4].

Interestingly, the most enriched DNA replication protein was the Pol *α* catalytic subunit (ranking 9) (Table 2; S2A Table). Pol *α* is recruited to the replication fork after the CMG complex (**<u>C</u>**dc45, **<u>M</u>**CM 2-7 subunits and the **<u>G</u>**INS complex) and activated by MCM10, triggering DNA unwinding at the origin of replication [76,77]. Pol *α* synthesizes RNA primers and physically interacts with RFC1 and RFA1 at the replication fork [75]. With the exception of PCNA and the MCM subunits, there are no functional studies for the core replication proteins. Some known replication factors did not display high scores and other known DNA replication proteins did not pass the filtering criteria including DNA polymerase epsilon catalytic subunit, MCM2, Replication Factor A (51 kDa subunit) and DNA topoisomerase II. This is possibly due to low copy numbers of these proteins at the replication fork or less efficient interaction with the DNA.

Several proteins of unknown function are likely to be DNA replication or repair proteins based on the presence of conserved domains. For example, Tb927.3.5370 and Tb927.9.10400 contain regions of similarity to type II DNA topoisomerases. DNA topoisomerases manage the topological state of the DNA, with type II enzymes catalyzing the passage of one double stranded DNA duplex through a break in another DNA duplex in an ATP dependent mechanism [78,79]. A range of processes including DNA replication, DNA repair, sister chromatid segregation, chromosome condensation and catenation rely on type II topoisomerases [80,81].

Another interesting candidate, Tb927.9.15070, has similarity to DNA processing A (DprA) superfamily (NCBI Conserved Domains Database, cl22881). DprA is a bacterial member of the larger, extremely diverse recombination mediator protein (RMP) family. RMPs facilitate binding of RecA-like recombinases (RecA, Rad51) to DNA damage sites for the repair of broken chromosomes and other types of DNA lesions [82].

To test if the DNA replication proteins identified are enriched on nascent DNA, we calculated the FC EdU/ThD chase for these proteins. We compared the ThD chase sample with the three EdU pulse replicates and the FC estimated in each EdU pulse was added to obtain an average FC. From this analysis, RFC4 and MCM4 were only detected in the EdU and absent in the ThD chase sample (INF value). The remaining known DNA replication proteins have a total FC ranging from 0.6 to 4.7, where PCNA is highly enriched (S2C Table) compared to the ThD chase sample. Therefore, DNA replication proteins such as MCM4 and PCNA are in close proximity to the replication fork in the EdU pulse and as the labeled DNA moves away from the fork in the thymidine chase, these proteins are not enriched or are no longer detected. Our results are in agreement with iPOND results from other systems [28,66,83],

### Other enriched proteins

Our data set revealed DNA repair as a GO term (S2D Table) with a fold enrichment of 5.43 (P-value 8.10 • 10^-8^). RAD54 (rank 12) and RAD51 (rank 378) participate in repairing DNA double-strand breaks by homologous recombination and have physical and functional interaction. RAD54 drives branch migration and stimulates RAD51 strand exchange activity [84] whereas RAD51 mediates homology search, strand invasion, and D-loop formation steps [85]. When RAD51 was knocked out in *T. brucei*, the parasites were more sensitive to DNA damaging agents [86]. The fact that we are enriching proteins related to DNA repair on nascent DNA might indicate that DNA repair is active during new synthesis or shortly after DNA is replicated.

Using SV40 minichromosomes, nucleosomes were shown to assemble immediately behind the DNA replication fork with the first deposited nucleosome detected at a distance of only ~250 bp from the replication fork [87]. An estimated 37 kbp of *T. brucei* DNA can be labeled in a 10 min EdU pulse. Therefore, if nucleosome assembly occurs similarly near the active fork, it would be expected to find enrichment of these assembly/remodeling proteins on nascent DNA in *T. brucei*. Accordingly, we found that chromatin organization had the highest fold enrichment (10.70; P-value of 2.91 • 10^-12^) (Fig 2A, S3 Table). Nucleosome components such as histone 3 (H3), 4 (H4), 2A (H2A), 2B (H2B), 2A variant Z (H2A.Z), 2B variant (H2BV) and 3 variant (H3V) were all detected in the data set (S2A and S2D Tables). H2A.Z and H2BV are specifically deposited at TSSs, whereas H3V was found at TTSs [19]. Histone chaperones such as FACT, contribute to rapid nucleosome assembly at the replication fork, and chromatin remodeling enzymes such as Isw1 help load and position nucleosomes during DNA replication [69]. Accordingly, there are representatives from at least 3 chromatin remodeling complexes present in the data set, namely both FACT subunits (rank 212, 375), two INO80 RuvB-like proteins (rank 338,397), and 3 of the 4 ISWI complex proteins (rank 96, 255, 324). In addition, several other putative nucleosome assembly proteins were detected (S2E Table). Interestingly, when comparing average FC EdU/ThD chase, several of the chromatin associated proteins, especially the FACT components, were enriched on nascent DNA further supporting the notion that nucleosome assembly occurs in proximity to the replication fork (S2E Table).

There are numerous examples replication and transcription responding to similar cues from the chromatin landscape in eukaryotes. Genome-wide studies in *Drosophila*-revealed that early replicating regions were associated with increased transcriptional activity, activating chromatin marks (acetylation), and increased ORC binding, while late replicating regions were associated with repressive histone marks (methylation) [88,89]. Genome-wide analyses of *T. brucei* ORC1 binding revealed surprisingly few origins of replication (~100) that map to the boundaries of their DGC revealing an unprecedented level of functional interaction between transcription and DNA replication [15]. Additionally, in the related trypanosomatid *L. major*, genome-wide studies indicated that initiation and timing of DNA replication depend on RNA pol II transcription dynamics that also serve as a driving force for nucleosomal organization [14].

In support of an intimate association between transcription and DNA replication machineries, the transcription GO term had a 9.91-fold enrichment (P-value of 3.66 • 10^4^) (Fig 2A; Table 3), and STRING analysis showed transcription proteins interacting with all clusters except for nuclear transport and protein metabolism (Fig 2B). Notably, the main subunits of all three nuclear RNA polymerases were identified which may reflect the facts that RNA pol I in trypanosomes transcribes the genes encoding the parasite’s major cell surface proteins [90] and that RNA pol III transcribes genes often located in convergent SSRs [19,91]. A small percentage (7%) of early replicating TbORC1 binding sites are associated with convergent SSRs, and these sites could overlap with RNA Pol III transcription start sites [15,19,91]. Other important transcription proteins identified include TATA-box binding protein (rank 16), two elongation factors (rank 98, 110), and several basal transcription initiation factors (rank 68, 72, 131, 147, 240). These findings are in accordance with enrichment of transcription machinery in iPOND studies conducted in other systems [33,66,83].

**Table 3.** List of Transcription related protein identified in *T. brucei* iPOND

| **Tb427 Protein ID** | **Tb927 Protein ID** | **Product Description** | **Mol. Weight [kDa]** | **Ranking Position** | **Total Score** |
| --- | --- | --- | --- | --- | --- |
| Tb427.04.5320 | Tb927.4.5320 | Component of IIS longevity pathway SMK-1, putative | 97 | 7 | 134.65 |
| Tb427.10.15950 | Tb927.10.15950 | TATA-box-binding protein | 29 | 16 | 109.87 |
| Tb427.10.2780 | Tb927.4.5020 | DNA-directed RNA polymerase II subunit RPB1 | 170 | 29 | 104.03 |
| Tb427tmp.47.0010 | Tb927.11.1390 | class I transcription factor A, subunit 1 | 53 | 68 | 78.98 |
| Tb427.01.1700 | Tb927.1.1700 | AATF protein, putative (Apoptosis-antagonizing transcription factor) | 55 | 69 | 78.31 |
| Tb427.02.5030 | Tb927.2.5030 | transcription initiation protein, putative | 80 | 72 | 77.59 |
| Tb427.01.1680 | Tb927.1.1680 | Transcription elongation factor 1 domain-containing protein | 26 | 98 | 73.27 |
| Tb427.04.3510 | Tb927.4.3490 | DNA-directed RNA polymerases II and III subunit RPB6, putative | 16 | 99 | 73.21 |
| Tb427.08.5090 | Tb927.8.5090 | DNA-directed RNA polymerase I largest subunit | 196 | 104 | 73.03 |
| Tb427tmp.03.0760 | Tb927.11.370 | repressor activator protein 1 | 93 | 105 | 72.94 |
| Tb427.02.3580 | Tb927.2.3580 | transcription elongation factor s-II, putative | 52 | 110 | 72.53 |
| Tb427.08.5980 | Tb927.8.5980 | TFIIH basal transcription factor complex helicase subunit, putative | 92 | 131 | 70.54 |
| Tb427tmp.01.5370 | Tb927.11.13810 | ATP- dependent RNA helicase, putative | 63 | 143 | 70.00 |
| Tb427tmp.01.1200 | Tb927.11.9430 | TFIIH basal transcription factor subunit | 41 | 147 | 69.28 |
| Tb427tmp.02.0970 | Tb927.11.3480 | Sin-like protein conserved region, putative | 79 | 200 | 45.94 |
| Tb427.01.540 | Tb927.1.540 | DNA-directed RNA polymerase III, putative | 127 | 217 | 42.92 |
| Tb427.10.8720 | Tb927.10.8720 | CCR4-NOT transcription complex subunit 10, putative | 61 | 236 | 41.14 |
| Tb427tmp.47.0008 | Tb927.11.1410 | class I transcription factor A, subunit 3 | 47 | 240 | 40.68 |
| Tb427.10.15370 | Tb927.10.15370 | DNA-directed RNA polymerases I and III subunit RPAC1, putative | 37 | 242 | 40.47 |
| Tb427tmp.160.4220 | Tb927.9.5710 | general transcription factor IIB | 38 | 245 | 40.30 |
| Tb427.03.1270 | Tb927.3.1270 | PRP38 family, putative | 61 | 256 | 39.67 |
| Tb427.04.1310 | Tb927.4.1310 | ZFP family member, putative zin finger transcription factor | 47 | 294 | 36.37 |
| Tb427tmp.03.0450 | Tb927.11.630 | RNA polymerase I second largest subunit | 179 | 323 | 16.03 |
| Tb427.04.3810 | Tb927.4.3810 | DNA-directed RNA polymerase II subunit 2, putative | 134 | 336 | 13.67 |
| Tb427tmp.211.3840 | Tb927.9.12900 | RNA polymerase-associated protein LEO1, putative | 65 | 354 | 10.92 |
| Tb05.5K5.70 | Tb927.5.4420 | nucleolar RNA helicase II, putative | 69 | 380 | 38.72 |
| Tb427.08.1510 | Tb927.8.1510 | ATP-dependent RNA helicase DBP2B, putative | 62 | 381 | 8.68 |
| Tb11.v5.0414 | Tb927.10.540 | ATP-dependent RNA helicase SUB2, putative | 49 | 407 | 4.28 |

Interestingly, proteins involved in pre-mRNA splicing were also enriched on nascent DNA (GO term fold enrichment of 7.03 P-value of 2.21 • 10^-7^) (Fig 2A, S2F Table). Proteins that are part of the spliceosome such as SmB (rank 42), SmD2 (rank 163) and U5-40K (rank 308) were identified. Also, proteins involved in polyadenylation and capping were identified (S2E Table). Recently, the link between nucleosome occupancy, RNA pol II levels and splicing elements in *T. brucei* was addressed [92]. In this study, RNA pol II sites were found in close proximity to regions marked by the histone variant H2A.Z and were associated with TSS. The RNA Pol II enrichment could indicate transcription pausing when encountering nucleosomes and can cause a recruitment of the splicing factors [92]. H2A.Z, RNA pol II subunits and splicing factors are enriched in our data set suggesting that co-transcriptional splicing events might occur in near proximity of newly replicated DNA.

### Analysis of Newly Identified Factors

To begin a functional analysis of our data set, we selected the putative replication protein RFC3 (Tb927.10.7990; rank 15) and a protein of unknown function, Tb927.3.5370 (rank 123) that contained a conserved Type II-like topoisomerase domain. To date, there have been no functional studies for any of the RFC subunits. Using TritrypDB resources, Tb7990 appeared to be important for parasite fitness in RIT-seq analysis but showed little change in transcript levels during the cell cycle [93,94]. In contrast, Tb5370 was not greatly affected in RIT-seq but showed cell cycle-dependent changes in transcript levels that peaked during S phase. In the high-throughput TrypTag study [47], both proteins localized to the nucleus.

Tb5370 was selected because of its strong expression during S-phase and its possible association with type II topoisomerases that relieve topological constraints produced in many processes including DNA replication and transcription. The NCBI Conserved Domain Database [52] revealed similarity to a Top2c domain (domain PTZ00108, E-value 5.16 x 10^-5^). However, protein alignment of Tb5370 with other type II topoisomerases showed Tb5370 was lacking key residues involved in topoisomerase activity. By inspecting for other conserved motifs in Tb5370 and the corresponding homologs in other trypanosomatids, we identified an FHA (forkheaded associated domain) phosphopeptide ligand motif and the cyclin dependent kinases (CDK) phosphorylation site motif (S3 Fig). Importantly, using the phosphoproteomic data available for *T. brucei* [95], three serine residues are phosphorylated in the CDK phosphorylation motif of Tb5370 suggesting it is likely controlled by a kinase (S3 Fig). CDKs phosphorylate protein substrates that are associated with regulation of cell cycle transitions [96], such as DNA synthesis and mitosis [97]. In addition, these kinases also phosphorylates proteins involved in DNA damage [98]. Since Tb5370 is predicted to have FHA motifs which are present in proteins involved in DNA repair and transcription [99], and to be phosphorylated by CDKs, we decided to further evaluate this gene to assess if is essential in *T. brucei*.

Tb7990 was annotated as RFC3. Replication factor C (RFC) is a five-subunit complex that catalyzes the ATP-dependent loading of PCNA onto DNA for replication and repair. The five essential subunits contain the ATP-binding Walker A motif with the consensus sequence GxxxxGKK, the magnesium ion-binding Walker B motif hhhhNExx that is required for ATP hydrolysis [100], and the SRC motif, also called an arginine finger, that senses bound ATP and participates in ATP hydrolysis [101]. RFC5 subunits differ slightly in having an imperfect Walker B motif that prevents ATP hydrolysis [101]. Tb7990 contains the three conserved motifs (S4 Fig), however Tb7990 has distinct substitutions in the Walker A (GxxxxGK<u>T</u>) and Walker B (hhhh<u>D</u>Exx) motifs.

To evaluate the nuclear localization of these proteins during the cell cycle, each factor was fused at the C-terminus with the PTP tag and expressed from their respective endogenous locus in clonal procyclic cell lines. To directly examine localization of the tagged proteins during DNA replication, we labeled bulk DNA with DAPI and newly synthesized DNA with EdU (Fig 3). We applied structured illumination microscopy (SIM) to visualize at high-resolution colocalization of the tagged protein with EdU foci following 10 min of labeling. Both proteins displayed nuclear localization throughout the cell cycle. The Tb5370-PTP signal localized more at the nuclear periphery reminiscent of nuclear pores and did not appear to overlap with EdU foci (Fig 3A). This distinct pattern was retained even during mitosis (data not shown). Tb7990-PTP localized as several punctate foci and was excluded from the nucleolus. While there was colocalization with EdU foci, the two patterns did not precisely overlap, suggesting Tb7990 may have roles other than DNA replication (Fig 3B). There are four different RFC complexes in eukaryotes: RFC1-RFC, Ctf18-RFC, Elg1-RFC and Rad17-RFC [102]. These three complexes share the four small RFC subunits (RFC2, 3, 4, and 5) but differ in the large subunit [103,104]. RFC1-RFC is the canonical RFC complex that acts as a processivity factor for DNA polymerases during replication [105]. Ctf18-RFC is involved in chromatid cohesion [106], Elg1-RFC plays role in genome stability [107] and Rad17-RFC is part of the DNA damage checkpoint response [108].

**Figure 3.**
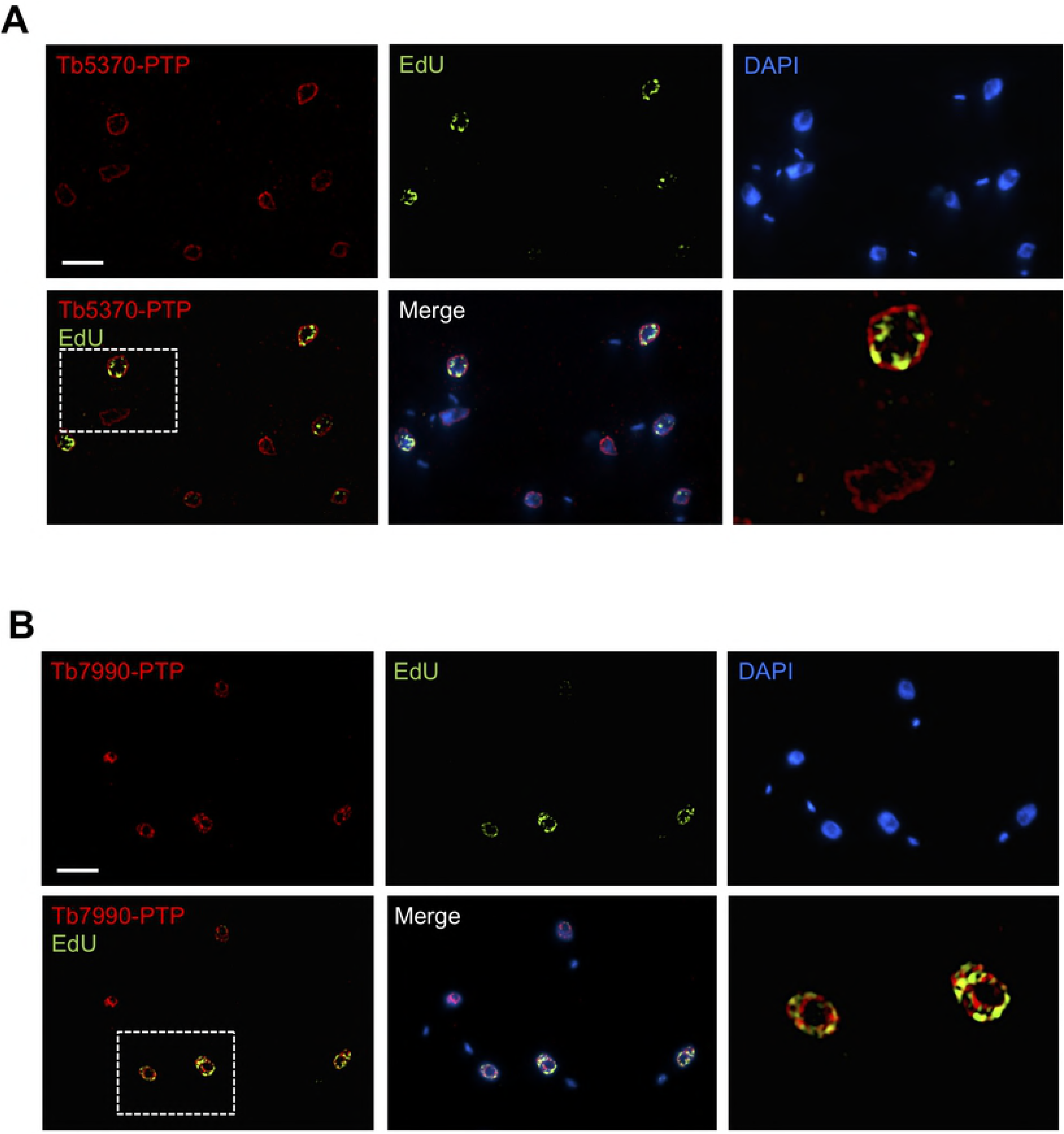
Intranuclear localization of two iPOND candidates. Localization of PTP-tagged proteins in asynchronous populations labeled with EdU using SIM microscopy. PTP tag was detected with anti-protein A (red), EdU incorporation (green) and DNA was stained with DAPI (blue). Enlargements correspond to the white dashed boxes. **(A)** Tb5370-PTP tagged cell line **(B)** Tb7990-PTP tagged cell line. Size bar, 5 μm.

The function of Tb5370 was further explored using a stemloop RNAi construct to deplete its mRNA in a tetracycline-dependent manner. However, there was no loss of fitness following induction of RNAi. Tb5370 RNAi cells before and after tetracycline induction were harvested for RNA isolation and mRNA levels were monitored using RT-qPCR (S5 Fig). *Tb5370* mRNA levels were reduced 80-90% in several clones that were analyzed. It appears that Tb5370 is not essential under standard growth conditions, however a role in DNA repair cannot be ruled out at this point.

To further investigate the function of Tb7990, this gene was silenced using inducible RNAi. We prepared total RNA from uninduced and tetracycline-treated cells after 48 hr of growth and monitored mRNA levels using RT-qPCR (Fig 4A, inset). *Tb7990* mRNA levels were reduced ~75% with a corresponding decrease in fitness starting on day 3 (Fig 4A). To characterize the effect of Tb7990 depletion on cell cycle progression and DNA replication directly, cells before and after tetracycline induction were examined using fluorescence microscopy. Since the kinetoplast (K) divides before the nucleus (N), the trypanosome cell cycle stage can be determined by examining DAPI-stained cells in an asynchronous population [56]. Additionally, the appearance of 2 mature basal bodies is used to mark the onset of kDNA replication (1N1K^∗^) and is easily visualized using the YL1/2 antibody that is a marker for mature basal bodies. The percentage of cells with 1N1K, 1N1K^∗^, 1N2K and 2N2K configurations were scored (Fig 4B, S6 Fig). Tb7990 silencing led to a transient increase in 1N1K^∗^ cells, but resulted in an overall decline after 4 days (from 40 to 18%). There was a gradual decline of 1N1K cells (from 42 to 28%) that was accompanied by an increase in 0N1K cells (zoids) from less than 1 to 20% of the population. Cells with other abnormal configurations also increased by day 5 (from 0 to 12%). Thus, these data indicate that silencing Tb7990 resulted in impaired cell cycle progression. It is important to note that in procyclic *T. brucei*, cell division can proceed in the absence of nuclear division or even nuclear S phase to give rise to anucleate cells (zoids) [109–111]. Additionally, the appearance of zoids and disruption of cell cycle progression was reported for silencing of the DNA replication initiation proteins TbORC1 and MCM subunits [4,40].

**Fig 4.**
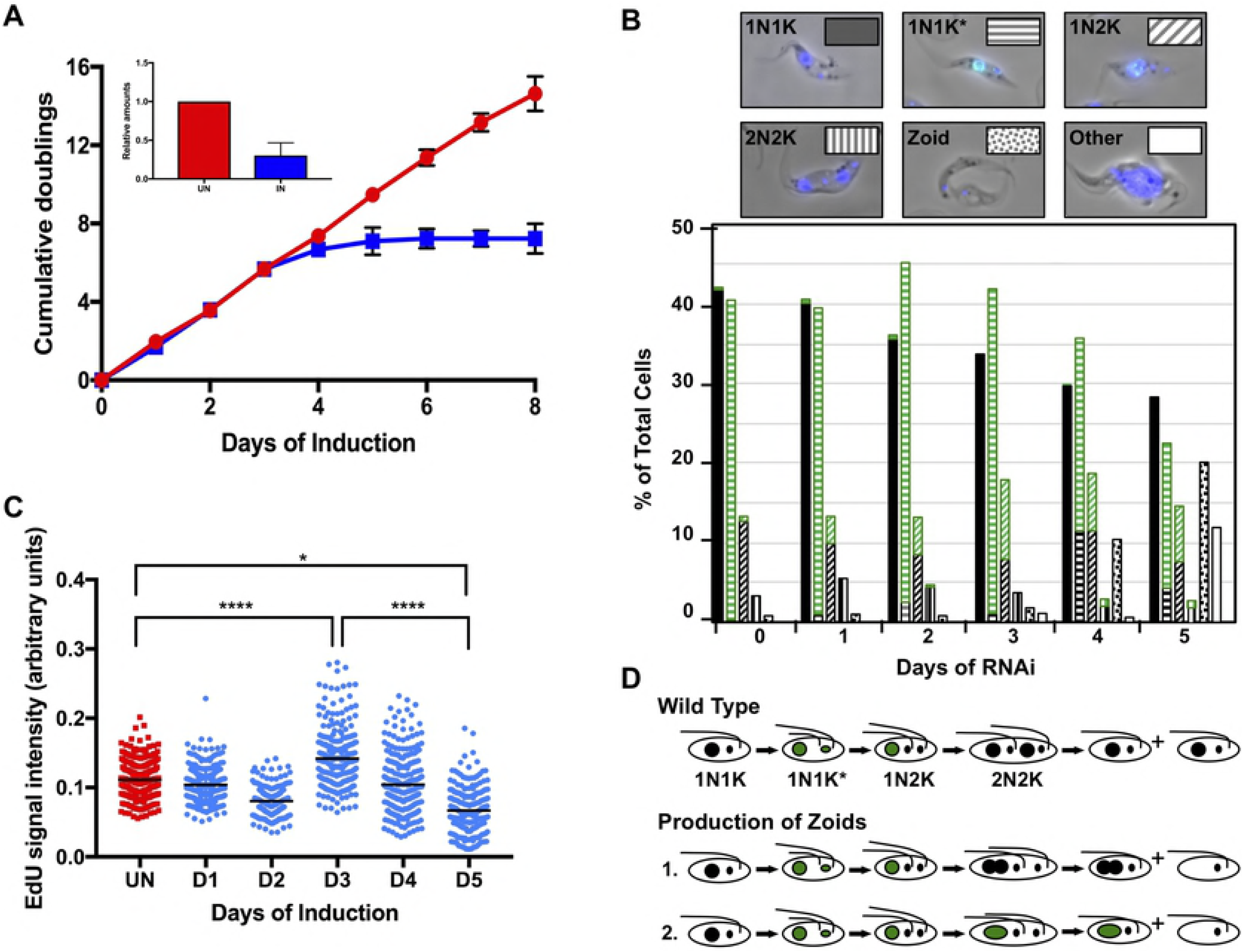
Silencing of Tb7990 by RNAi. **(A)** Tb7990 Growth curves, in the presence and absence of tetracycline. Inset, qPCR **(B)** Quantification of cell cycle stages following Tb7990 RNAi. Cells were induced for RNAi (0 – 5 days), EdU labeled for 10 min, fixed, and processed for detection of EdU, basal bodies and DNA. Cells (~325/day) were scored by fluorescence microscopy according to the number of kinetoplasts (K) and nuclei (N), EdU+ nuclei, and number of basal bodies. Some categories (e.g. 2N0K, 2N1K, multiple nuclei) comprised <2% of the total and grouped as Other. Images show examples of each cell type scored and key to bar graphs. **(C)** Each dot represents the mean EdU signal per nucleus for each condition after a 30 min pulse with 150 μM EdU. **(D)** Model for production of zoids (adapted from [112]). Larger circles, nuclei; small ovals, kDNA; black, EdU-; green, EdU+.

To directly test whether Tb7990-depleted cells were defective in DNA replication, the same cells were metabolically labeled with EdU following the induction of Tb7990 dsRNA, and nascent DNA was detected by indirect immunofluorescence microscopy. Cells were scored for the absence (EdU-) or presence (EdU+) of EdU labeling. In an asynchronous uninduced population, 41.1% of the cells exhibited an EdU-dependent fluorescence signal (EdU+) with the great majority of cells being 1N1K^∗^ (40.2%). Following Tb7990 depletion, there was an initial increase in the percentage of EdU+ cells that subsequently declined to 26% at Day 5. The percentage of EdU+ 1N1K^∗^ cells decreased while the percentage of EdU+ 1N2K cells increased indicating that progression through S phase was impaired and cells were not progressing through mitosis to G2 (Fig 4B, S6 Fig).

In addition to the overall decline in EdU+ cells, the fluorescence intensity of the EdU+ cell population also decreased following 5 days of Tb7990 silencing (S6 Fig). To evaluate statistical significance, we used CellProfiler [43] to score fluorescence intensity in at least 220 EdU+ cells for each time point (Fig 4C). In uninduced cells, there is a distribution of fluorescence intensity ranging from 0.055-0.20 representing cells at the various phases of DNA replication in an unsynchronized population. Interestingly, there was an increase in fluorescence intensity observed at Day 3 in which the mean intensity increased and the range was broader (0.064-0.28) compared to uninduced (P-value <0.0001). During this time point some of the highest EdU fluorescence intensities were recorded and this timepoint corresponded to an increase in 1N2K cells as well as the percentage of 1N2K cells that were EdU+. By the last day of the RNAi induction, the mean EdU fluorescence intensity decreased even below uninduced values and had the lowest intensities as well as the lowest range (0.011-0.18) compared to all days (D3/D5 P-value <0.0001; Un/D5 P-value <0.02). Together these data demonstrate a role in DNA replication by both a reduction in the number of EdU+ cells and decreases in EdU incorporation.

The Tb7990 RNAi DNA replication defect was similar to the defects reported for silencing *T. brucei* ORC1, MCM subunits (3, 5 and 7) and PCNA in procyclic cells [4,25,40]. Knockdowns of these genes led to a gradual decline in the number of 1N1K cells that was accompanied by a gradual increase in zoid cells (0N1K) and a subtle increase in the 1N2K population, although there was variation in the percentages of cells at each of these cell stages. The accumulation of 1N2K cells can be the result of incomplete mitosis (a nuclear segregation defect) or some perturbation of S phase preventing the cells from initiating mitosis. Both defects can result in progeny where one cell receives some proportion of nuclear material while the other cell receives only a kinetoplast (zoid)(Fig 4D) due to the lack of a classical mitosis to cytokinesis checkpoint in procyclic trypanosomes [109–111]. The combined accumulation of zoids and transient increase in 1N2K cells appears to be a hallmark of DNA replication defects.

## Conclusion

Here, we have established the iPOND technology, for the first time in a parasitic system. It is a powerful tool that provided a global view of proteins that are enriched at nascent DNA under unperturbed replication conditions in *T. brucei* PCF cells. Our data suggest that DNA replication, transcription, chromatin organization and pre-mRNA splicing events all occur on and or near nascent DNA. These different cellular processes may be coordinated or just occur in the vicinity of each other. Based on our observations, we propose nucleosomes are assembled close to the replication fork followed by RNA pol II recruitment, transcription, and co-transcriptional RNA processing. Further studies are needed to determine how these processes are linked and co-regulated, and how rapidly they are initiated during DNA replication.

In addition, our data set has provided a list of proteins of unknown function that can be characterized to determine their function in DNA replication and/or at nascent DNA. They could represent essential trypanosomatid-specific factors or extremely divergent homologues of known replication factors and, therefore, represent promising targets of chemotherapeutic intervention. The initial characterization of the protein candidate Tb7990, a replication factor C subunit, indicated an essential role in DNA replication that was quantified using EdU incorporation and microscopy assays.

However, many of the DNA replication proteins identified in our data set have not been studied and their future characterization using our strategy to detect replication defects will contribute to the understanding of DNA replication in *T. brucei*.

## Acknowledgements

This work was partially supported by a UMass Graduate School Grant to MCR, and a Faculty Research Grant to MMK. NIH grant AI126455 to MMK supported this work. We thank David Cortez and Jared Nordman for discussions and ideas. We are grateful to Christian Janzen for providing the H3 antibodies, Paul Englund for the mtHsp70 antibodies, and Bibo Li for the pKOORC1-Hyg plasmid. The superresolution microscopy data were gathered in the Light Microscopy Facility Nikon center of Excellence at the Institute for Applied Life Sciences, UMass Amherst, with support from the Massachusetts Life Sciences center.

